# The reticulon homology domain of Pex30 generates membrane curvature at ER subdomains for lipid droplet biogenesis

**DOI:** 10.64898/2026.04.08.717014

**Authors:** Morgan House, Nikhil Nambiar, Steve M Abel, Amit S Joshi

**Affiliations:** Department of Biochemistry & Cellular and Molecular Biology, University of Tennessee, Knoxville 37916; Department of Chemical and Biomolecular Engineering, University of Tennessee, Knoxville 37916

## Abstract

Lipid droplets (LDs) are dynamic organelles that store neutral lipids and form in the endoplasmic reticulum (ER) membrane. Formation of new LDs is a controlled process and requires proteins with specific functions to form and grow from the ER membrane without any defect. *In vitro* studies have suggested a role for membrane curvature in LD emergence from the ER. Here, we use the membrane-shaping protein Pex30 to investigate the impact of ER membrane curvature on LD biogenesis and morphology. We modified the reticulon homology domain (RHD) of Pex30, which is responsible for tubulating the ER membrane, by extending the short hairpin transmembrane domains (TMD). The Pex30 (TMD) mutants cannot tubulate the ER membrane and generate less local membrane curvature that WT Pex30. Additionally, these mutants are unable to restore delayed LD biogenesis observed in cells devoid of Pex30. Our results indicate that Pex30 RHD generates local membrane curvature at ER subdomains that drives formation of new LDs.

## Introduction

Lipid droplets (LDs) are unique storage organelles that emerge *de novo* from the endoplasmic reticulum (ER). The core of a LD contains neutral lipids such as triacylglycerols (TAG) and sterol esters (SE), that are encapsulated by a phospholipid monolayer. Nascent LD biogenesis begins with the accumulation of neutral lipids within the ER bilayer, called LD nucleation, to form a lens-like structure (Choudhary *et al*., 2015). Once these neutral lipids reach a critical concentration of 5-10 mol %, they bud into the cytosol to form a LD (Walther *et al*., 2017; Olzmann and Carvalho, 2019). In yeast, LDs stay connected to the ER membrane (Jacquier *et al*., 2011). While this process begins with neutral lipid synthesis, proteins localized to these early sites of LD biogenesis have been shown to facilitate LD biogenesis. These ER membrane proteins include Sei1, Ldb16 and their interacting proteins Ldo16 and Ldo45, Pex30, Yft2, Pah1, Spo7, and Nem1 in yeasts and seipin, MCTP1/2, FIT2, GPAT4, and LDAF1 in mammalian cells (Fei *et al*., 2008; Adeyo *et al*., 2011; Wang *et al*., 2014, 2018; Choudhary *et al*., 2015, 2018, 2020; Grippa *et al*., 2015; Eisenberg-Bord *et al*., 2018; Joshi *et al*., 2018, 2021; Teixeira *et al*., 2018; Chung *et al*., 2019; Chen *et al*., 2021; Klug *et al*., 2021). The roles of these proteins in nascent LD biogenesis varies from binding TAG and initiating LD nucleation in the ER bilayer to determining the direction of LD budding (Chorlay *et al*., 2019). Another factor that influences nascent LD biogenesis is ER membrane curvature. *In vitro,* studies showed that LDs preferably assemble in ER tubules with high membrane curvature and reduced surface tension (Ben M’barek *et al*., 2017; Santinho *et al*., 2020). The critical TAG concentration required for LD formation is lower in curved ER tubules than in flat ER membrane sheets. Thus, LD nucleation can be achieved by increase of membrane curvature (Santinho *et al*., 2020). However, very little is known about how ER membrane curvature affects LD biogenesis *in vivo*.

Some studies have shown that overexpression of ER-shaping proteins such as reticulons in yeast and mammals, and Yop1 in *S. cerevisiae*, and cells devoid of Atlastin in *C. elegans* leads to smaller LDs, providing evidence that altering global ER shape can promote LD emergence (Klemm *et al*., 2013; Niemelä *et al*., 2025). On the other hand, depletion of reticulon and reticulon-like protein, Yop1, in *S. cerevisiae* does not affect LD morphology or abundance (Joshi *et al*., 2018). In *C.* elegans, it was shown that depletion of ER tubule-forming proteins along with FIT2 leads to a decrease in the number and size of LDs (Chen *et al*., 2021; Apte and Joshi, 2022). These studies suggest a role for membrane curvature in reducing the surface tension required for neutral lipids to coalesce into a mature LD (Ben M’barek *et al*., 2017; Santinho *et al*., 2020). However, these proteins are canonical ER-shaping proteins, and likely to affect LD biogenesis through changing overall ER shape. Are there proteins that could affect local ER membrane curvature at the LD biogenesis sites to directly affect formation of new LDs? Here we investigate the role of Pex30, a reticulon-like ER membrane protein enriched at ER subdomains that are sites for nascent LD formation (Joshi *et al*., 2018; House *et al*., 2025). We demonstrate the role of Pex30 reticulon homology domain (RHD), which has been shown to tubulate ER, in LD biogenesis (Joshi *et al*., 2016).

Pex30, an ER-resident protein, has two short hairpin transmembrane domains (TMD) which are part of the reticulon homology domain (RHD) responsible for tubulating ER membrane (Joshi *et al*., 2016). Thus, the Pex30 RHD is similar to reticulons and reticulon-like proteins. In addition, Pex30 has a dysferlin (DysF) domain and a domain of unknown function (DUF) (Vizeacoumar *et al*., 2004; Ferreira and Carvalho, 2021). Pex30 is part of a family of proteins, Pex28, Pex29, Pex31, and Pex32, that affect peroxisome size and abundance but are not essential for peroxisome function (Vizeacoumar *et al*., 2003, 2004; Mast *et al*., 2016; Deori *et al*., 2023). Pex30 is localized to the ER membrane where it forms several puncta in a wild type (WT) cell (Joshi *et al*., 2016). While some of these sites could be utilized for peroxisome vesicle formation, others are used as sites for formation of new LDs (Joshi *et al*., 2016, 2018; Wang *et al*., 2018). Interestingly, some of these Pex30 puncta are shared as both peroxisomes and LDs associated with it (Joshi *et al*., 2018). While Pex30 RHD is required for ER membrane shaping, the DysF domain is able to bind phosphatidic acid (PA), a precursor of diacylglycerol (DAG) and TAG (Joshi *et al*., 2016; Ferreira *et al*., 2025; House *et al*., 2025). We have shown that Pex30 is recruited to the ER subdomains enriched with PA via DysF domain in *sei1Δ*. Pex30 also has been found to form complexes with Pex30-like proteins Pex28, Pex29, Pex31, and Pex32 at different ER-organelle membrane contact sites (Ferreira and Carvalho, 2021). Additionally, lack of Pex30 results in smaller and clustered LDs with ER membrane proliferation around them, a delay of LD biogenesis and defect in targeting of proteins such as Dga1 to the LDs (Joshi *et al*., 2018). Even though Pex30 is at the LD formation sites, whether the Pex30 RHD shapes the ER membrane at these subdomains and affects LD biogenesis remains unknown.

In this study, we investigate the role of Pex30 RHD in generating membrane curvature for LD biogenesis. We mutated the two short membrane hairpins, TMD1 and TMD2 of the RHD, adding hydrophobic amino acids so that the hairpins can more easily span the ER bilayer rather than being predominantly in the cytosolic side of the ER bilayer. We use these mutant forms of Pex30 to investigate the ability of Pex30 to shape the ER membrane *in vivo* and *in silico*. We find that the Pex30 TMD mutants when overexpressed are unable to restore ER shape in the reticulon mutant and generate less local membrane curvature compared to WT Pex30. Furthermore, we investigate the effects of these TMD mutants on LD biogenesis and morphology. These mutants are unable to rescue the delay in formation of new LDs as observed in *pex30Δ*. In addition, TMD mutants affect the accumulation of Pex30 in *sei1Δ* mutant and are unable to restore defective LD morphology observed in *sei1pex30Δ*, indicating that the RHD of Pex30 is essential for efficient LD formation and biogenesis. Together, these results provide evidence that Pex30 RHD generates local ER membrane curvature, potentially lowering the energy barrier necessary for facilitating LD emergence from the ER.

## Results and Discussion

### Pex30 RHD is required for localization to the tubular ER

Previously, it was demonstrated that extending the short hairpin TMD length of human reticulon-4 affected its localization and function (Zurek *et al*., 2011). Using a similar approach, we extended the length of Pex30 TMD1, TMD2 or both TMD1 and TMD2 (TMD1.TMD2) by adding 10 hydrophobic residues including 3 acidic residues in the middle of the short hairpin TMD. This extension enables the TMD to span both leaflets of the ER bilayer (Figure 1A and 1B). Reticulons and reticulon-like proteins including Pex30 localize to regions of high membrane curvature such as ER tubules and edges of the ER sheets but do not localize to perinuclear ER. Pex30 exhibits punctate structure on the nuclear envelope (NE) possibly localizing at highly curved nuclear pores (Joshi *et al*., 2016). We tested if altering the TMD length changed the localization of Pex30. To do this, we tagged all the Pex30 TMD mutants and WT Pex30 with GFP at the C-terminus. We found that GFP tagged Pex30 and Pex30 TMD mutants were expressed and targeted to the ER membrane (Figure 1C). As expected, Pex30-GFP did not localize to perinuclear ER. In contrast, we found that Pex30 (TMD1), Pex30 (TMD2) and Pex30 (TMD1.TMD2) localized to NE suggesting that Pex30 short hairpin TMDs are required to exclusively localize the protein to high membrane curvature regions (Figure 1C). Interestingly, Pex30 (TMD2) still exhibited some cells with no NE localization. Our data shows that lengthening of the Pex30 short hairpin TMDs, either one or both, reduces the ability of Pex30 to exclusively localize to high curvature regions.

**Figure 1.**
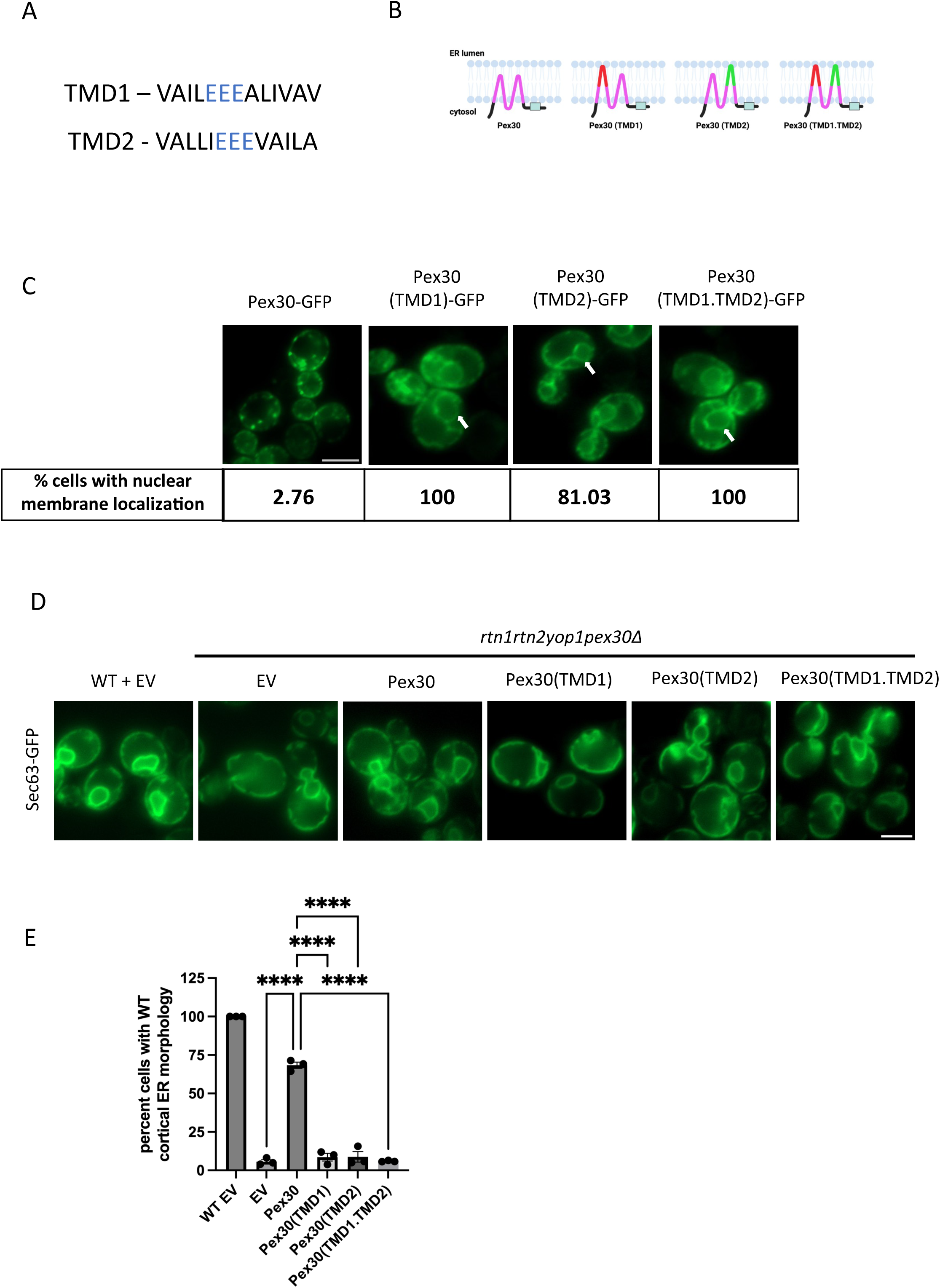
Topology and function of Pex30 TMD mutants. A. Amino acids added to hairpin regions of Pex30 to generate Pex30(TMD1) and Pex30(TMD2). Blue amino acids are acidic glutamate residues added to extend the short hairpin to bi-pass the ER bilayer. B. Cartoon depicting the predicted orientation of WT Pex30, Pex30(TMD1), Pex30(TMD2), and Pex30(TMD1.TMD2) in the ER membrane. C. Widefield images (WF) of WT cells expressing Pex30-GFP and GFP tagged Pex30 TMD mutants on a plasmid expressed under Pex30 promoter. White arrows denote Pex30 TMD mutants localized to the nuclear membrane (ER sheets). >100 cells per genotype were analyzed for nuclear membrane localization, and the percent cells showing this phenotype is shown in the table. Bar = 4 µm. D. WF images of *rtn1rtn2yop1pex30Δ* expressing Sec63-GFP as ER marker and WT Pex30 and Pex30 TMD mutants under *RTN1* promoter. Cells were imaged in logarithmic phase. Bar = 4µm. EV; empty vector. E. Quantification of images in D showing the percentage of cells with WT-like cortical ER morphology. Bars show the mean from three independent experiments and standard error of the mean (SEM). >100 cells per genotype from each replicate were analyzed and compared using ordinary one-way ANOVA and Dunnett’s multiple comparison test. (****, p < 0.0001)

### Pex30 RHD is required for tubulating the ER membrane

In *S. cerevisiae*, reticulons and reticulon-like proteins such as Rtn1, Rtn2, and Yop1 mainly regulate the ER shape. Deletion of all three proteins leads to more ER sheets than ER tubules suggesting they contribute to tubulating the ER membrane (De Craene *et al*., 2006; Voeltz *et al*., 2006). Cells devoid of Pex30 do not exhibit change in cellular ER morphology as it is a less abundant protein and localized to ER subdomains. Previously, we demonstrated that overexpression of Pex30 under a strong *RTN1* promoter leads to restoration of ER shape in the *rtn1rtn2yop1Δ* mutant (Joshi *et al*., 2016). In this study, we investigate if the Pex30 TMD mutants are able to tubulate the ER membrane. To do that, we overexpressed untagged Pex30, Pex30 (TMD1), Pex30 (TMD2), and Pex30 (TMD1.TMD2) under a *RTN1* promoter in *rtn1rtn2yop1pex30Δ* cells labelled with Sec63-GFP. As expected, we found that Pex30 restored the ER shape as a significant number of cells exhibited wild type (WT)-like cortical reticulated ER membrane. However, overexpression of Pex30 (TMD1), Pex30 (TMD2) and Pex30 (TMD1.TMD2) did not restore the ER morphology (Figure 1D and 1E). Our results suggested that short hairpin TMDs are essential for Pex30 to tubulate the ER membrane.

Next, we evaluate the ultrastructure of the ER membrane using transmission electron microscopy (TEM). WT cells show reticulated cortical ER (Figure 2A) whereas *rtn1rtn2yop1pex30Δ* cells exhibit continuous and wavy cortical ER membrane (Figure 2B). Overexpression of Pex30 but not the Pex30 (TMD1), Pex30 (TMD2), or Pex30 (TMD1.TMD2) under the *RTN1* promoter was able to rescue the ER shape to WT-like, which is consistent with fluorescence microscopy data (Figure 2C-2F). Our data indicates that the short hairpin TMDs are important for Pex30 localization and ER shaping function.

**Figure 2.**
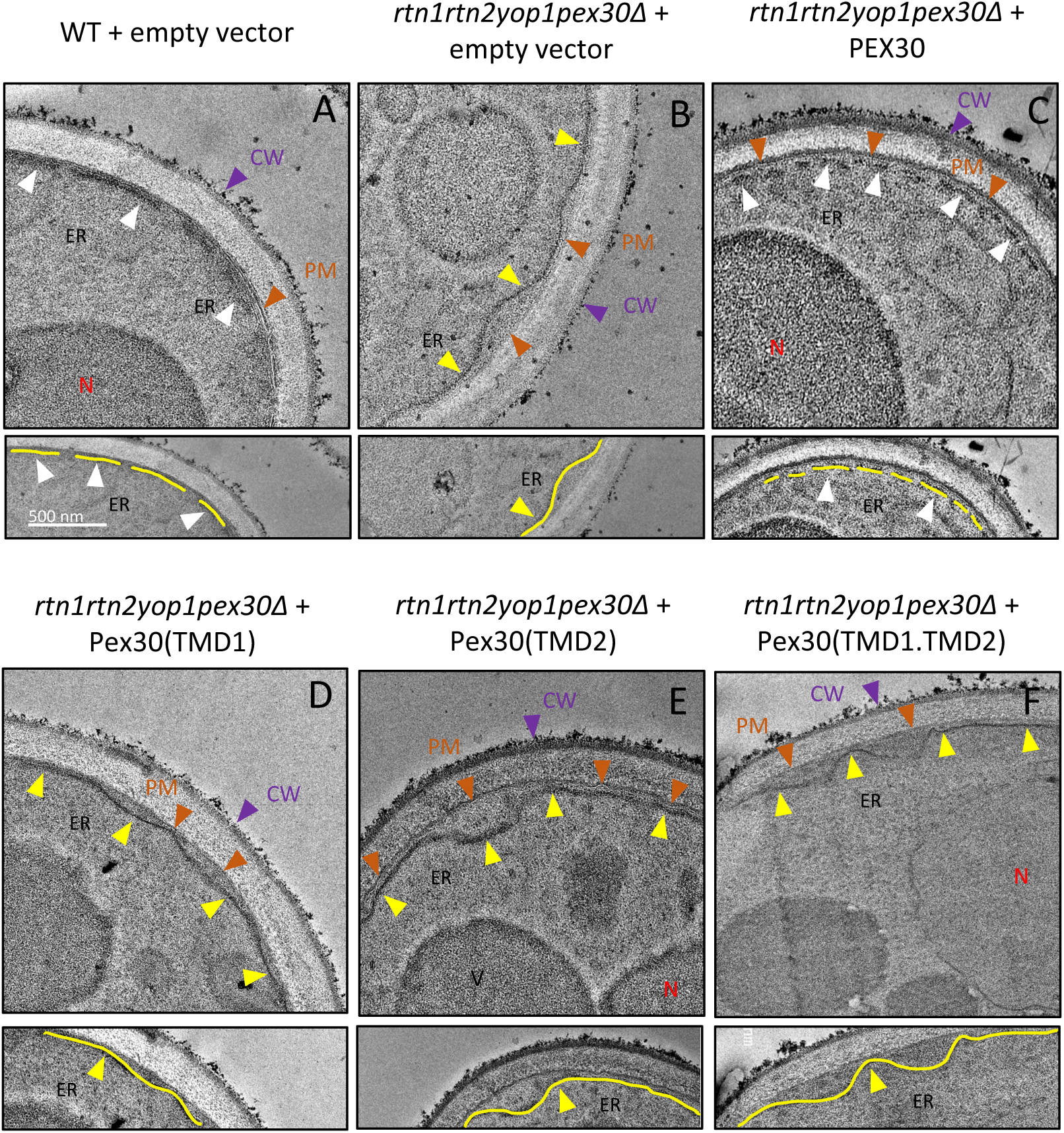
Pex30 TMD mutants are unable to tubulate the ER membrane. A-F. Electron microscopy images of the indicated strains expressing empty vector (EV) or Pex30 and Pex30 TMD mutants under *RTN1* promoter on a plasmid. White arrowheads denote ER tubules. Yellow arrowheads denote ER sheets. Purple arrowheads denote the cell wall (CW). Orange arrowheads denote the plasma membrane (PM). N is nucleus and V is vacuole. Bottom panels are zoomed in images of top panel with annotations of ER membrane in yellow.

### Pex30 RHD is required to maintain ER membrane curvature and thickness at single molecule level

To probe the molecular-scale impact of the Pex30 RHD on a membrane, we performed coarse-grained molecular dynamics (MD) simulations of the RHD, with and without TMD inserts, in a model ER membrane. Representative snapshots show the two short hairpin TMDs embedded in the lipid bilayer and spanning from the cytosolic leaflet to the lumenal leaflet (Figure 3A). Visual inspection suggested that the RHDs locally deform the membrane. To quantify the apparent deformation, we calculated the local mean curvature in the region surrounding the protein (Figure 3B). Time-resolved analysis revealed persistent positive curvature near the RHD. Figure 3C shows the average value of the mean curvature within 1 nm of each RHD variant over time. The wildtype Pex30 RHD generated the largest mean curvature. The Pex30 (TMD2) insert alone resulted in a modest reduction, whereas the Pex30 (TMD1) insert generated a more pronounced decrease, both alone and in combination with the Pex30 (TMD2) insert. The Pex30 (TMD1.TMD2) double insert exhibited the lowest mean curvature near the protein. We further examined membrane thickness in the vicinity of the RHD (Figure 3D). The WT RHD exhibited bilayer thickening near the RHD. The Pex30 (TMD2) alone produced a similar thickness profile with slightly less thickening of the bilayer. Introducing the Pex30 (TMD1) insert resulted in thinning of the membrane near the protein, both alone and with the Pex30 (TMD2) insert. Thus, the MD simulations show that modifying the first short hairpin results in a more pronounced reduction in local membrane curvature and thickness than modifying the second short hairpin, which has only a modest effect.

**Figure 3.**
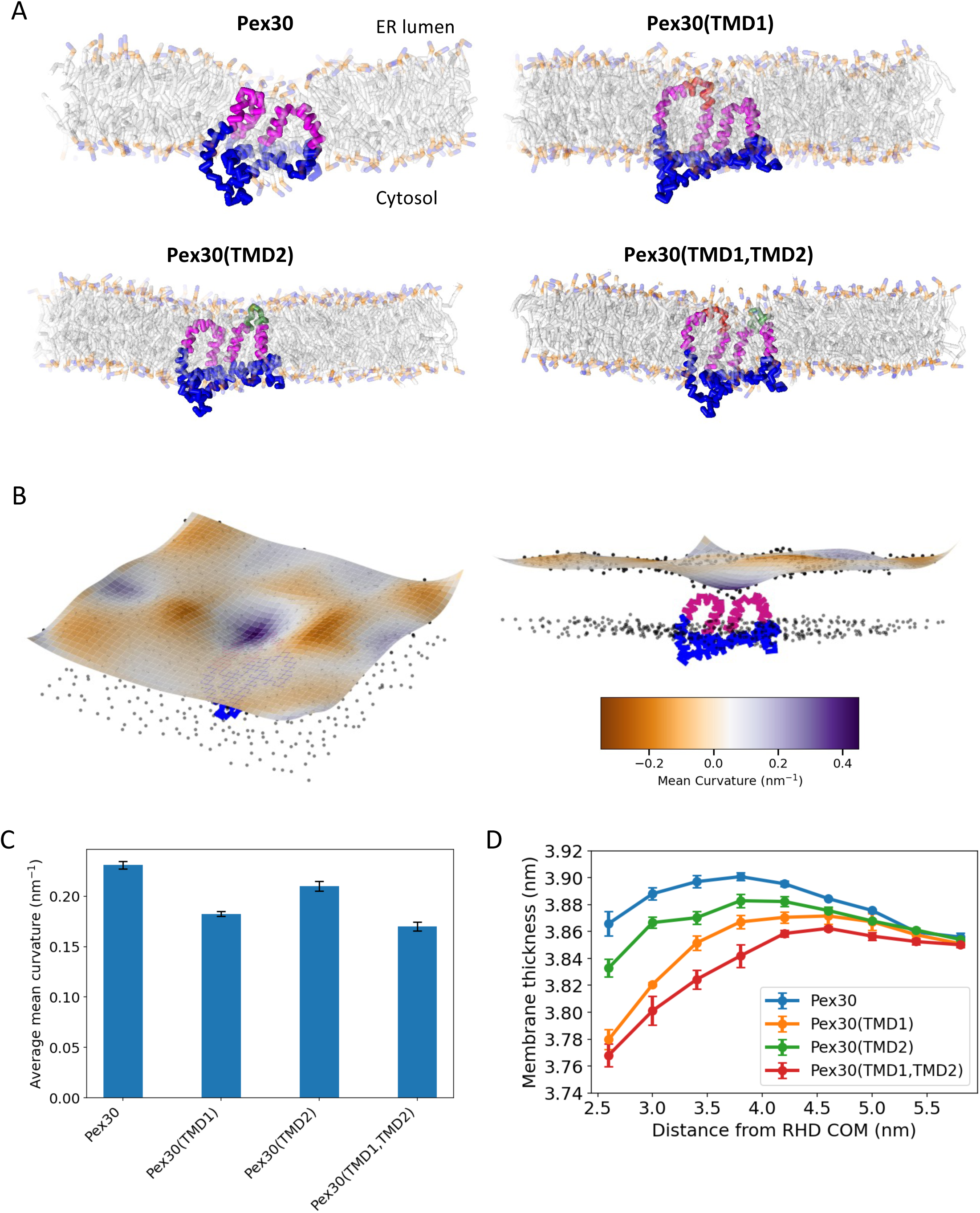
Coarse-grained MD simulations to characterize the Pex30 RHD and its mutants in a lipid bilayer. A. Snapshots from simulations of the Pex30 RHD in a model ER bilayer. The transmembrane domains of the RHD are shown in magenta. TMD1 and TMD2 insertions are shown in red and green, respectively. B. Surface representation of the lumenal membrane surface. Black points denote the positions of phosphate beads from lipid headgroups. For each simulation frame, a smooth bivariate spline was fit to the phosphate beads of lipids in the lumenal leaflet and used to determine the local mean curvature. C. Mean curvature of the lumenal surface around the RHD. The average mean curvature was calculated within a region extending 1 nm from the center of mass of the RHD over 1 µs of simulation time. Error bars represent the standard error from three independent simulation trajectories. D. Thickness of the membrane around the RHD. The mean thickness was calculated in annular regions as a function of distance from the center of mass of the RHD.

### Pex30 RHD is required to restore the LD morphology in *sei1pex30Δ*

Next, we tested functionality of Pex30 TMD mutants using the *sei1pex30Δ* mutant. Previously, we showed that *sei1pex30Δ* exhibits a severe growth defect and defective LD morphology consisting of enlarged as well as small, and clustered LDs (Joshi *et al*., 2018; Wang *et al*., 2018). When we overexpressed Pex30-GFP in *sei1pex30Δ* cells, it rescued the growth defect and accumulated at ER-LD contact sites as in *sei1Δ* (Joshi *et al*., 2018; Wang *et al*., 2018; Ferreira *et al*., 2025; House *et al*., 2025). These cells exhibited fewer and enlarged LDs similar to *sei1Δ*. Expression of truncated Pex30-GFP devoid of RHD or DysF domains were unable to rescue the growth defect and restore the LD phenotype (House *et al*., 2025). When Pex30 (TMD1)-GFP and Pex30 (TMD1.TMD2)-GFP mutants are overexpressed in *sei1pex30Δ*, cells still retain the LD phenotype as in *sei1pex30Δ* cells and these Pex30 TMD mutants do not accumulate into large puncta at ER-LD contact sites as WT Pex30-GFP (Figure 4A-4D). Additionally, Pex30 (TMD1)-GFP and Pex30 (TMD1.TMD2)-GFP mutants are unable to rescue the growth defect of *sei1pex30Δ* at 37°C (Figure 4E). These results suggests that membrane curvature induced by first short hairpin is important for Pex30-GFP accumulation and rescue of *sei1pex30Δ* growth defect.

**Figure 4.**
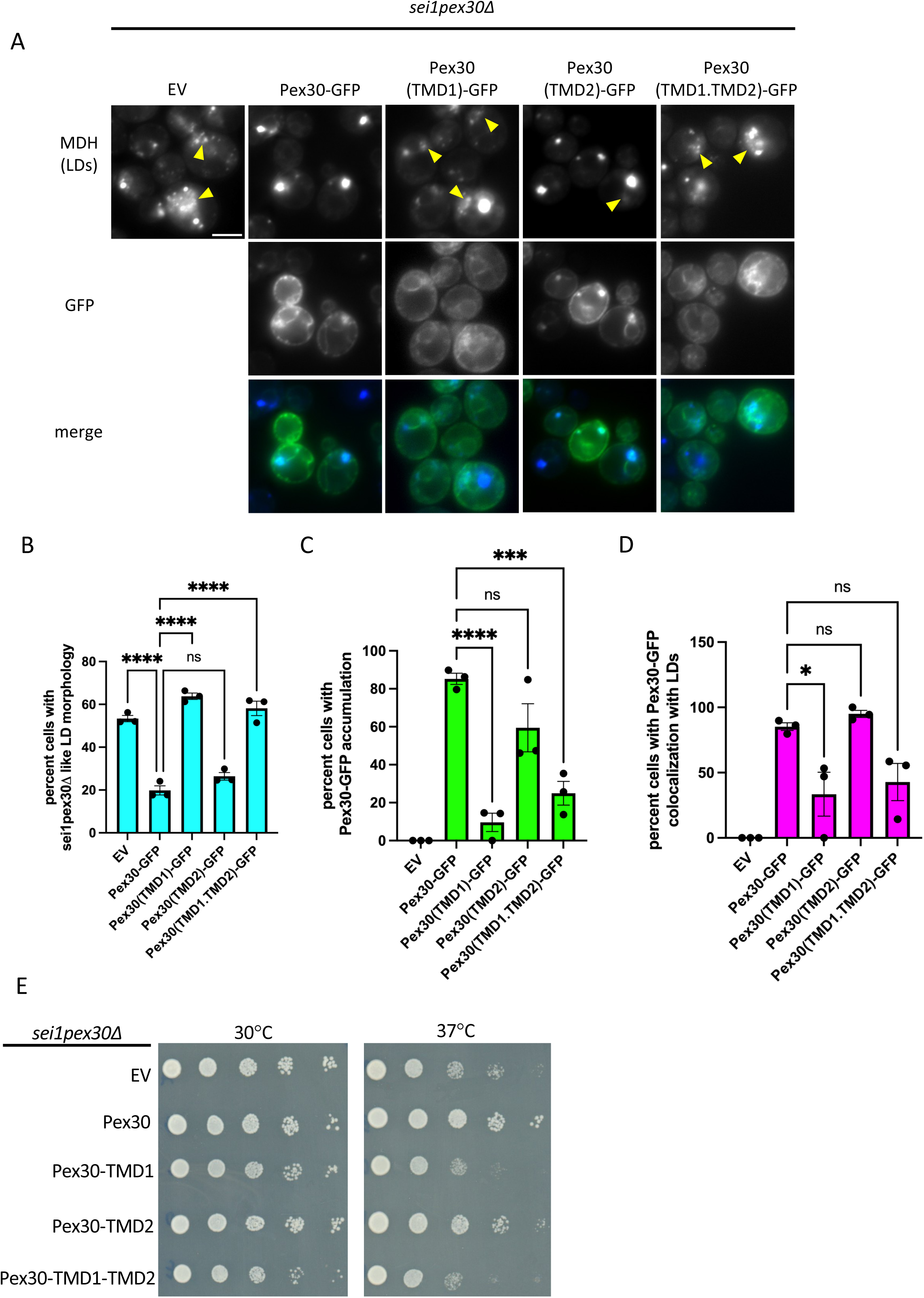
Pex30 RHD is required to restore LD morphology of *sei1pex30*Δ. A. WF images of *sei1pex30*Δ expressing EV or GFP-tagged Pex30 and Pex30 TMD mutants on a plasmid. Cells were imaged in logarithmic phase and stained with monodansylpentane (MDH) to visualize lipid droplets (LDs). Bar = 4µm. B-D. Quantification of images in A showing, (B) the percentage of cells with *sei1pex30*Δ like LD phenotype, (C) the percentage of cells with Pex30-GFP accumulation, (D) the percentage of cells with Pex30-GFP colocalization with LDs. Bars show the mean from three independent experiments and standard error of the mean (SEM). >100 cells per genotype from each replicate were analyzed and compared using ordinary one-way ANOVA and Dunnett’s multiple comparison test. (*, p < 0.05; ns – not significant, **, p < 0.01; ***, p < 0.001; ****, p < 0.0001) E. Spot test with tenfold serial dilutions of *sei1pex30*Δ cells expressing EV or GFP tagged Pex30 and Pex30 TMD mutants on a plasmid shown in A were spotted on SC media without leucine and incubated for two days at 30°C and 37°C.

The Pex30 (TMD2)-GFP mutant exhibits fewer and enlarged LDs similar to *sei1Δ*, suggesting it is functional like WT Pex30-GFP control (Figure 4A and 4B). Furthermore, Pex30 (TMD2)-GFP does accumulate into large puncta at ER-LD contact sites as in Pex30-GFP (Figure 4A, 4C and 4D) and rescues the growth defect of *sei1pex30Δ* at 37°C (Figure 4E). Together these results suggest that Pex30 (TMD2) can maintain function of WT Pex30. Consistent with the MD simulation findings, our results indicate that the first short hairpin of Pex30 is essential to retain Pex30 function at the ER subdomains for membrane curvature and LD biogenesis.

### Pex30 RHD is required for formation of new LDs

Next, we checked if Pex30 TMDs play a role in formation of new LDs. Previously, we reported that loss of Pex30 leads to delay in the induction of new LDs (Joshi *et al*., 2018). Therefore, we determined if the Pex30 (TMD1) and Pex30 (TMD2) mutants are able to rescue the delay in LD formation. To do this, we overexpressed Pex30-GFP, Pex30 (TMD1)-GFP, Pex30 (TMD2)-GFP and Pex30 (TMD1.TMD2)-GFP in *are1are2dga1pex30Δ* with *LRO1* expressed under *GAL1* promoter and labelled with Erg6-mCherry. LDs were labelled with the monodansylpentane (MDH) dye. We find that expression of all Pex30 TMD mutants were unable to rescue the delay in LD formation. Moreover, unlike the cells devoid of Pex30, cells expressing Pex30 TMD mutants exhibit a dominant negative effect on LD biogenesis as they show decreased LD formation at 5-hour timepoint (Figure 5A). These results suggest that Pex30 short hairpins generate membrane curvature that is important for formation of new LDs.

**Figure 5.**
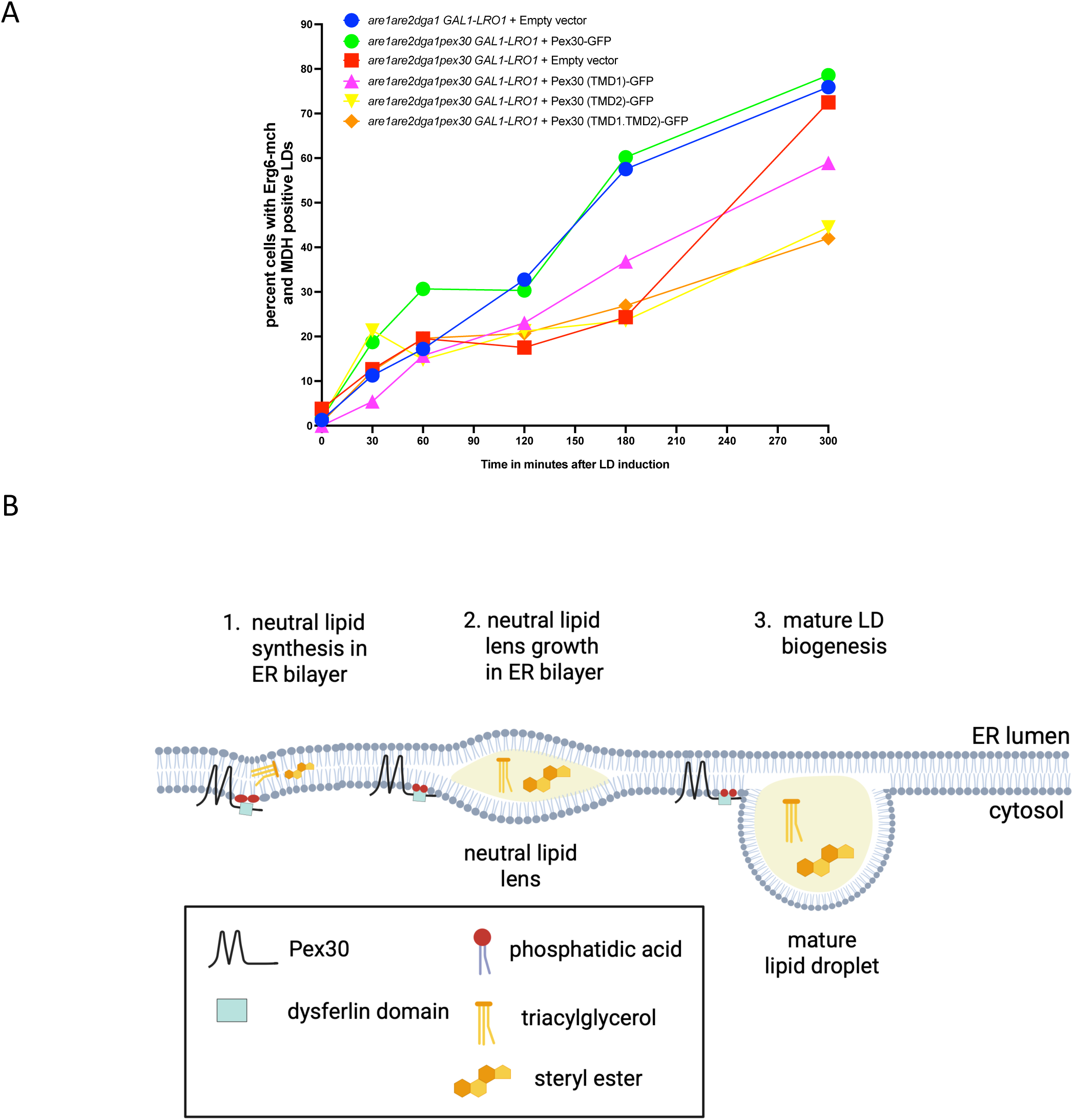
Pex30 TMD mutants display delayed LD biogenesis. A. LD induction experiment showing the percentage of cells with Erg6 and MDH positive LDs over time. Three independent experiments were performed, and >100 cells per genotype from each replicate were analyzed. B. Model for LD biogenesis in *S. cerevisiae*.

### Pex30 short hairpin TMDs generate membrane curvature at ER subdomains for LD biogenesis

In this report, we demonstrate that Pex30 generates membrane curvature at ER subdomains through its RHD consisting of two short hairpin TMDs that affect LD biogenesis. We alter the Pex30 RHD by extending the two short hairpin TMDs. We show that Pex30 RHD is necessary to target Pex30 to tubular ER. Additionally, Pex30 TMD mutants, unlike Pex30, are unable to restore ER morphology in *rtn1rtn2yop1pex30Δ* (Figure 1 D and E, Figure 2) which is consistent with the model that Pex30 RHD is required for ER membrane tubulation (Joshi *et al*., 2016). Furthermore, molecular dynamics (MD) simulations show that a single molecule of Pex30 generates membrane curvature around it in a model ER bilayer, and that extending its TMDs reduces the local membrane curvature and thickness (Figure 3 B and C). MD simulations provide evidence that Pex30 generates local curvature at ER subdomains and affects ER membrane thickness around the protein through its RHD.

Recently, studies showed that Pex30 DysF domain binds phosphatidic acid (PA) (Ferreira *et al*., 2025; House *et al*., 2025). We propose a model where Pex30 is recruited through its DysF domain to early LD biogenesis sites by binding PA (Figure 5B). PA is a precursor for diacylglycerol (DAG) which is converted to the neutral lipid triacylglycerol (TAG) that is stored in LDs. Thus, we propose that PA recruits Pex30 at ER subdomains where LDs form prior to the nucleation step (Figure 5B). How does Pex30 facilitate LD biogenesis at these sites? In this study, we investigate how membrane curvature generated by Pex30 RHD enables formation of new LDs in the ER membrane. Our findings show that the Pex30 RHD is responsible for generating membrane curvature and increasing membrane thickness. We propose that Pex30 generates local membrane curvature and increases membrane thickness thus facilitating accumulation of neutral lipids (Figure 5B). As the lens grows within the ER bilayer, the local membrane curvature generated by Pex30 lowers the energy barrier needed for LD nucleation while also reducing the surface tension in the ER for LD growth (Figure 5B). Further studies could focus on identifying the magnitude of curvature and type of phospholipids that could be enriched at these sites and whether Pex30 forms oligomers, similar to other RHD-containing ER shaping proteins, which would enhance membrane shaping at these sites (Shibata *et al*., 2008; Fuentes *et al*., 2023; Xiang *et al*., 2023). Additionally, LD biogenesis proteins may not be recruited properly to ER subdomains without Pex30-generated membrane curvature, leading to a delay in the formation of new LDs (Choudhary *et al*., 2020). Identifying proteins and lipids enriched at Pex30 subdomains would shed light on mechanistic details of LD biogenesis.

## Materials and Methods

### Yeast strains and plasmids

All yeast strains, plasmids, and primers used in this study are listed in supplemental table 1. Chromosomal gene deletion and fluorescence tagging of genes was performed using PCR-based targeted homologous recombination and tetrad dissection (Longtine, et al., 1998).

Plasmids used in this study were generated by double restriction enzyme digestion of the plasmid at either the BamHI and SalI restriction sites or the EcoRI and HindIII restriction sites. Pex30 TMD mutants were generated using PCR. Amino acid insertions were incorporated by adding their respective nucleotide sequences to primers and amplifying regions of Pex30. Each PCR product contains homologous overhangs, and these products and digested plasmid were transformed into yeast cells using standard lithium acetate transformation methods. Yeast and *E.coli* plasmids were isolated with D2004 and D4016 kit respectivley (Zymo Research). Yeast plasmids were amplified in competent *E. coli* cells (New England Biolabs C3030H). Pex30 TMD mutant plasmids were confirmed through PCR and sequencing (Plasmidsaurus).

### Yeast media and growth conditions

Yeast cells were grown in YPD (1% yeast extract, 2% peptone, 2% glucose) or synthetic complete (SC) media with respective amino acid dropouts for plasmid selection (0.67% yeast nitrogen base without amino acids (USBiological), amino acid mix (USBiological), and 2% glucose). For LD induction experiments, SC without leucine was prepared with 2% raffinose for precultures and 2% galactose for LD induction.

### Lipid droplet induction experiment

For LD induction, cells were precultured in SC media with 2% raffinose. Cells were pelleted, and fresh SC media with 2% galactose was added to cells along with monodansylpentane (MDH) (Abcepta) dye at a concentration of 1mM. Cells were grown at 30°C in the dark and imaged at 30 minutes, 1 hour, 2 hours, 3 hours, and 5 hours post induction. For time point 0, cells precultured in raffinose were incubated with MDH dye for 30 minutes in the dark at 30°C and imaged. Cells were quantified based on the colocalization of MDH and Erg6 puncta.

### Fluorescence microscopy

For fluorescence microscopy, cells were precultured overnight, diluted to 0.2-0.3 OD_600_ units and imaged in mid-log phase. LDs were visualized by staining cells with MDH dye for 30 minutes at 30°C in the dark. All images in this study are widefield images obtained on an inverted Zeiss 900/Airyscan laser scanning confocal microscope equipped with Colibri 7-channel solid-state fluorescence light source with two filter sets for widefield microscopy and diode lasers and gallium arsenide phosphide. Images were acquired using a 63×/1.4 NA objective lens.

### Image quantification and analysis

All images in this study were analyzed and quantified using ImageJ software. Cells were quantified manually using Z-stacked images. Percentage values are the cells showing a specific phenotype divided by the total number of cells analyzed. All statistics in this study were performed using GraphPad Prism Version 10.6.1, and the specific analyses conducted are indicated in the figure legends.

### Electron microscopy

Sample preparation for EM was done as described previously (Choudhary *et al*., 2015). Yeast cells were cultured overnight in SC media and diluted to 0.1 OD_600_ in 15mL. Cells were grown until 1.0 OD_600_, and 15 OD units of cells were pelleted. Cells were resuspended in 15mL of fixative buffer (1% glutaraldehyde (Electron Microscopy Sciences) and 0.2% paraformaldehyde (Electron Microscopy Sciences) in 40mM potassium phosphate buffer, pH 7.0). Cells were rotated in fixative buffer for 1 hour at RT and shifted to 4°C overnight with rotation. Next, cells were pelleted and washed twice with 0.9% NaCl solution and once with water. Cells were incubated in a 2% solution of potassium permanganate for 5 min at RT. Cells were pelleted, resuspended in fresh 2% potassium permanganate, and rotated for 45 min at RT. Cells were pelleted and washed twice with water. Cells were incubated in a graded ethanol series of 30%, 50%, 70%, 90%, and 100% for dehydration. Cells were incubated in each ethanol solution for 10 min at RT with rotation and pelleted between each solution. Cells were stored in fresh 100% ethanol overnight at RT with rotation. The next day, cells were pelleted and rotated in a 2:1 ratio of 100% ethanol:Spurr’s resin for 4 hours at RT. Cells were pelleted and resuspended in a 1:1 ratio of ethanol:resin and rotated overnight at RT. The next day, cells were pelleted and resuspended in 1:2 ethanol:resin. Cells were rotated at RT for 4 hours and pelleted. Next, cells were resuspended in 100% resin and rotated overnight at RT. The next day, cells were pelleted and resuspended in fresh 100% resin. Cells were placed in BEEM capsule molds and pelleted. Molds were placed under vacuum for 4-5 hours to remove bubbles and transferred to 60°C for 3-4 days for resin polymerization.

Embedded samples were trimmed using glass knives, and ultrathin sections (70–100 nm) were collected using the Leica UC7 ultramicrotome (Leica Microsystems, Wetzlar, Germany). Sections were collected on 200-mesh carbon-coated copper grids (Electron Microscopy Sciences, Hatfield, PA, USA). The sections were stained with 2% uranyl acetate for 30 minutes, followed by 2% lead citrate for 5 minutes. Imaging was performed using a JEOL JEM-1400 FLASH transmission electron microscope (JEOL Ltd, Peabody, MA, USA) at 120 kV by using OneView Gatan camera (Gatan, Inc., Pleasanton, California, USA).

### Molecular dynamics simulation

Coarse-grained molecular dynamics simulations using the Martini force field were used to study individual, membrane-embedded RHDs (Periole and Marrink, 2013). We used the CHARMM-GUI Martini Maker to set up a model ER lipid bilayer composed of DOPC (65%), DOPE (18.5%), POPS (8.25%), DOPA (5.5%), and DOPS (2.75%) lipids (Jo *et al*., 2008; Qi *et al*., 2015). In total, the bilayer consisted of 800 lipids. The structure of the RHD was predicted using ColabFold on the full sequences of Pex30, Pex30(TMD1), Pex30 (TMD2), and Pex30 (TMD1.TMD2) (Mirdita *et al*., 2022). The RHD region of each was used in the simulations, which consisted of residues 60-254 in Pex30 without TMD inserts, and the same region plus additional residues for the TMD-insert variants. The resulting RHDs were then inserted into the bilayer using CHARMM-GUI. The PPM (positioning of proteins in membranes) server was used to determine the initial placement of the RHDs (Lomize *et al*., 2012). All simulations were performed in GROMACS 2021 using the Martini 2 force field (Van Der Spoel *et al*., 2005; Abraham *et al*., 2015). The LINCS algorithm was used for bonded interactions. Lennard-Jones and Coulombic interactions were cut off at 1.1 nm using the potential-shift-Verlet modifier and reaction field method respectively. Coulombic interactions were also cut off at 1.1 nm using the reaction field method. The temperature was set at 303.15 K, ensuring all lipids were in the liquid-disordered state, and a concentration of 0.15 mM NaCl was used to neutralize the charge of the system. The bilayer was equilibrated using the standard multi-step equilibration procedure from CHARMM-GUI. The equilibration consisted of four stages, where the restraints on the bilayer head groups were relaxed and the simulation time step was increased from 2 fs to 20 fs. During the equilibration process, temperature coupling was handled by the velocity-rescaling thermostat, and we used isotropic Berendsen pressure coupling with a reference pressure of 1 bar and compressibility of 3 × 10^−4^ bar^−1^ (Berendsen *et al*., 1984). In production runs, we used the velocity-rescaling thermostat and the Parrinello-Rahman barostat (Parrinello and Rahman, 1981; Bussi *et al*., 2007). The production run was simulated for 2 µs and the last 1 µs was used for analysis. Trajectories generated with GROMACS were analyzed in Python using the MDAnalysis package. PyMOL was used to create snapshots of the simulations(Michaud-Agrawal *et al*., 2011). Three independent trajectories were run for the Pex30 RHD and each of the mutants.

Membrane curvature around the protein was quantified using a surface-fitting approach, with Martini phosphate beads used to define the location of the bilayer surface. A smooth surface, 𝑧(𝑥, 𝑦), was fitted to the positions of the phosphate beads in the lumenal leaflet using a bivariate spline with a smoothing parameter of 3 × 𝑁, where 𝑁 was the number of phosphate beads in the leaflet. Using the smooth bivariate spline, first and second order derivatives of 𝑧 with respect to 𝑥 and 𝑦 (𝑧*_x_*, 𝑧*_y_*, 𝑧*_xx_*, 𝑧*_yy_*, 𝑧*_xy_*) were numerically calculated on a 2D grid with a spacing of 0.05 nm. The mean curvature at each point on the grid was calculated using

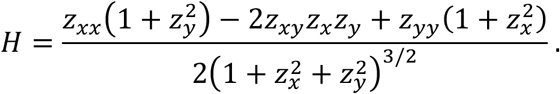

The mean curvature at the grid points within a 1-nm radius around the RHD’s center of mass (COM) was averaged to obtain the average mean curvature around the protein in a single simulation frame (Giblin, 2001). For each trajectory, the average mean curvature in this region was averaged over 1000 frames, sampled every 1 ns. We analyzed the lumenal side of the membrane because the RHD displaced lipids on the cytosolic side, making it challenging to reliably fit a surface near the COM of the RHD.

To calculate the membrane thickness around the RHD, phosphate bead positions were analyzed in successive annular regions, centered on the RHD’s COM, of 0.4 nm thickness in the 𝑥 − 𝑦 plane. Within each annular region, the membrane thickness was defined as the distance between the average 𝑧 position of the phosphate beads in the lumenal and cytoplasmic leaflets. The thickness is reported only in regions where lipid headgroups were consistently present in both leaflets.

## Acknowledgement

We thank Dr. William Prinz for reading the manuscript. Research reported in this publication was supported by NIH award R35 GM147189 and startup funds from University of Tennessee at Knoxville to ASJ and by the National Science Foundation (CBET-2217777) to SMA. MH was supported by NIH T32 award GM142621. We thank Jaydeep Kolape and Dr. Jennifer Rybak from Advanced Microscopy and Imaging Center (AMIC) at University of Tennessee in Knoxville for help with sample preparation and TEM imaging. Computer simulations were performed on the University of Tennessee Infrastructure for Scientific Applications and Advanced Computing (ISAAC) computational resources.

