## Supplemental Table 1 for "The reticulon homology domain of Pex30 generates membrane curvature at ER subdomains for lipid droplet biogenesis"

**Supplemental table 1: List of yeast strains, plasmids and primers used in this study.**

| **Yeast Strain List** | | | |
|---|---|---|---|
| **Strain number** | **Name** | **Genotype** | **Source** |
| AJY1000 | BY4741 | *MATa his3Δ1 leu2Δ0 met15Δ0 ura3Δ0* | Dharmacon Inc |
| AJY1315 | *rtn1rtn2yop1pex30Δ* | BY474? *MAT? his3Δ1 leu2Δ0 met15Δ0 ura3Δ0 rtn1::KanMx6 rtn2::KanMx6 yop1::KanMx6 pex30::HIS3* | Laboratory collection |
| AJY1069 | *sei1pex30Δ* | BY474? *MAT? his3Δ1 leu2Δ0 met15Δ0 ura3Δ0 sei1::KanMx6 pex30::KanMx6* | Laboratory collection |
| AJY1374 | *are1are2dga1Δ ERG6*-mCherry *GAL1-LRO1* | BY474? *MAT? his3Δ1 leu2Δ0 met15Δ0 ura3Δ0 are1::KanMx6 are2::KanMx6 trp1::URA3 dga1::Lox-HIS-Lox TRP1-GAL1-LRO1 ERG6-mCherry-HIS3* | Laboratory collection |
| AJY1375 | *are1are2dga1pex30Δ ERG6*-mCherry *GAL1-LRO1* | BY474? *MAT? his3Δ1 leu2Δ0 met15Δ0 ura3Δ0 are1::KanMx6 are2::KanMx6 trp1::URA3 dga1::Lox-HIS-Lox TRP1-GAL1-LRO1 ERG6-mCherry-HIS3 pex30::CaURA3* | Laboratory collection |
| **Plasmid list** | | | |
| **Plasmid number** | **Name** | **Description** | **Source** |
| pAJ1063 | Yeplac181-*PEX30*-GFP | GFP fused to C-terminus of *PEX30* in Yeplac181 backbone; Leu2 and ampicillin selection markers | Laboratory collection |
| pAJ1121 | Yeplac181- *PEX30* (TMD1)-GFP | GFP fused to C-terminus of *PEX30* TMD1 mutant in Yeplac181 backbone; Leu2 and ampicillin selection markers | This study |
| pAJ1122 | Yeplac181- *PEX30* (TMD2)-GFP | GFP fused to C-terminus of *PEX30* TMD2 mutant in Yeplac181 backbone; Leu2 and ampicillin selection markers | This study |
| pAJ1123 | Yeplac181- *PEX30* (TMD1.TMD2)-GFP | GFP fused to C-terminus of *PEX30* TMD1.TMD2 mutant in Yeplac181 backbone; Leu2 and ampicillin selection markers | This study |

| | | | |
|---|---|---|---|
| pAJ1064 | Yeplac181-proRTN1-*PEX30* | *PEX30* expressed under *RTN1* promoter in Yeplac181 backbone; Leu2 and ampicillin selection markers | Laboratory collection |
| pAJ1124 | Yeplac181-proRTN1-*PEX30*(TMD1) | *PEX30* TMD1 mutant expressed under *RTN1* promoter in Yeplac181 backbone; Leu2 and ampicillin selection markers | This study |
| pAJ1125 | Yeplac181-proRTN1-*PEX30*(TMD2) | *PEX30* TMD2 mutant expressed under *RTN1* promoter in Yeplac181 backbone; Leu2 and ampicillin selection markers | This study |
| pAJ1487 | Yeplac181-proRTN1-*PEX30*(TMD1.TMD2) | *PEX30* TMD1.TMD2 mutant expressed under *RTN1* promoter in Yeplac181 backbone; Leu2 and ampicillin selection markers | This study |
| pAJ1052 | Sec63-GFP | GFP fused to C-terminus of *SEC63* in Ycplac33 backbone; Ura3 and ampicillin selection markers | Laboratory collection |

**Primer list**

| **Primer** | **Sequence** |
|---|---|
| first insert-Pex30 (109-115 aa)- R | caccgccacaatcagcgcttcttcttccagaatcgccacATACTCAACGGTAGTAATAAA |
| first insert-Pex30 (116-122 aa) - F | gtggcgattctggaagaagaagcgctgattgtggcggtgTTTGAAACGCTGGTGAAGTAC |
| first insert-Pex30 (196-202 aa) - R | cgccagaatcgccacttcttcttcaatcagcagcgccacAAGTAGCCAGGTTATCATCAC |
| second insert-Pex30 (203-209 aa) - F | gtggcgctgctgattgaagaagaagtggcgattctggcgTTACCACCGAGAAGCTTGATG |
| Yep181-PEX30-600 F | ggataacaatttcacacaggaaacagctatgaccatgattTTCAGTACCAGCTCATTGTC |
| Yep181-PEX30+300 R | agggttttcccagtcacgacgttgtaaaacgacggccagtAATACTTTCCCATCCGCATA |
| Yep181-proRTN1-PEX30-F | ggataacaatttcacacaggaaacagctatgaccatgattTCTGTACGTGTGTGTGGACA |

| | |
|---|---|
| Yep181-gPEX30GFP-R | agggttttcccagtcacgacgttgtaaaacgacggccagtTATTACCCTGTTATCCCTAG |
| **Pex30**-proRTN1 R | TGGCTCTAGTCTCATGCACGTTAGTTGTGTTACCACTCATATTTGCGTGTGTGAATATAT |
| RTN1-Pex30 F | CAAGCGTATATATATATAATATATATTCACACACGCAAATATGAGTGGTAACACAACTAA |
